# The human hippocampus is involved in implicit motor learning

**DOI:** 10.1101/2024.08.30.610548

**Authors:** Guillermina Griffa, Alvaro Deleglise, Agustín Solano, Florencia Jacobacci, Gabriela De Pino, Valeria Della-Maggiore

## Abstract

Recent evidence suggests that the human hippocampus, traditionally associated with declarative memory, plays a role in motor sequence learning (MSL). However, the classic MSL paradigm depends initially on declarative learning. Therefore, it is critical to discern whether the participation of the hippocampus relates to its canonical role or to processing a general aspect of learning that transcends the declarative/non-declarative distinction. To address this issue, here we turn to visuomotor adaptation -a type of motor learning involving skill recalibration-which unlike MSL can be easily manipulated to eliminate the explicit component. Here, we examined the broader involvement of the hippocampus in procedural motor learning by using diffusion MRI to indirectly assess structural plasticity associated with memory consolidation in visuomotor adaptation (VMA) and an implicit-only version (IVMA). We found that both VMA and IVMA engaged the left posterior hippocampus in a learning-specific manner. Remarkably, while VMA induced only transient hippocampal alterations, IVMA elicited structural changes that persisted overnight, underscoring the reliance on implicit learning for enduring neuroplasticity. As expected, training on both visuomotor tasks impacted the microstructure of the cerebellum, the motor and the posterior parietal cortex. Notably, the temporal dynamics of changes in these regions closely paralleled those of the left hippocampus, suggesting that motor and limbic regions operate in a coordinated manner as part of the same neural network. Collectively, our findings support an active role of the hippocampus in procedural motor memory and argue for a unified function in memory encoding regardless of the declarative or non-declarative nature of the task.

## INTRODUCTION

In the 1950s, Brenda Milner’s extensive work on patient H.M. revealed that the surgical bilateral resection of the hippocampus markedly impaired the encoding of declarative memory (Scoville and Milner, 1957). Nevertheless, H.M. maintained the capacity to learn non-declarative motor tasks (Milner, 2005). Milner’s seminal work introduced the concept of specialized memory systems in the brain, which inevitably initiated a dichotomy in cognitive neuroscience, with episodic and semantic learning viewed as hippocampus-dependent and procedural learning as hippocampus-independent.

Recent neuropsychological and neuroimaging studies have challenged this traditional view, prompting a reassessment of the hippocampus’s role in procedural memory (Schendan et al., 2003; Fletcher et al., 2005; Gheysen et al., 2010; Albouy et al., 2013; Albouy et al., 2015; Deleglise et al., 2023). Using the motor sequence learning (MSL) paradigm involving executing a five-item sequence with the fingers of the non-dominant hand (Karni et al., 1995; Doyon et al., 2002), studies have shown that hippocampal dysfunction in amnesic patients significantly impairs both the rate of learning and memory consolidation (Döhring et al., 2017; Schapiro et al., 2019). Similar outcomes have been observed in epileptic patients with substantial hippocampal volume loss tested on the same task (Long et al., 2018). Thus, although hippocampal dysfunction may not hinder motor learning to the same extent as episodic learning, it has a measurable impact on the efficacy of memory encoding and consolidation.

In line with the neuropsychological evidence, we have recently demonstrated that MSL improvements in performance occurring during the rest periods interleaved with practice (micro-offline gains) are associated with an increase in hippocampal activity (Jacobacci et al., 2020a). This finding is reminiscent of neural replay observed during memory reactivation (Foster, 2017; Buzsáki, 2015; Buch et al., 2021). Functional changes were followed by rapid (∼30 min) alterations in hippocampal microstructure as evidenced by a reduction in mean diffusivity (MD), a metric derived from the diffusion tensor model (DTI) (Basser et al., 1994). Pioneering studies in rodents using DTI in combination with histology have shown that decreases in MD during learning are associated with synaptogenesis and astroglial expansion, two well-established markers of structural plasticity linked to LTP-like processes (Holtmaat and Svoboda, 2009; Sagi et al., 2012; Hofstetter and Assaf, 2017). Thus, we have proposed that the same neuronal ensembles that reactivate during the quiet rest periods of MSL training may undergo structural plasticity (Jacobacci et al., 2020a), a mechanism in line with the modern definition of an engram (Josselyn and Tonegawa, 2020).

Collectively, these studies suggest that, beyond its established role in declarative memory, the human hippocampus may also be relevant to procedural motor learning. However, the classic MSL paradigm depends initially on declarative learning as participants must first memorize the sequence. Thus, the participation of the hippocampus may be linked to its canonical role in processing the declarative component of the task (e.g. rehearsal of the sequence in working memory). A significant limitation of the MSL task is that it can not be manipulated to eliminate the contribution of explicit learning without substantially altering the paradigm (Krakauer et al., 2019). Alternatively, visuomotor adaptation (VMA) –a type of learning involving skill recalibration in response to visual perturbations– can be easily modified to isolate implicit learning (Taylor et al., 2014; Morehead et al., 2017; Albert et al., 2022). This makes it an ideal experimental paradigm for studying the broader involvement of the human hippocampus in procedural memory.

In this study, we aimed to determine whether the human hippocampus is involved in implicit motor learning. Building on our previous longitudinal study (Jacobacci et al., 2020a) we used MD, an indirect marker of structural plasticity, to quantify changes in microstructure induced by visuomotor adaptation in the short term (30 min) and the long term (24 h). To assess the hippocampus’s role in implicit motor learning, we examined how removing the explicit component of the classic VMA paradigm impacted on hippocampal microstructure using an implicit-only version (IVMA) (Morehead et al., 2017; Kim et al., 2018). We predicted that if the hippocampus is indeed required for procedural motor learning, both VMA and IVMA would induce structural changes in this structure. Additionally, based on the competition hypothesis, which suggests that implicit and explicit learning compete for error in visuomotor adaptation (Albert et al., 2022), we hypothesized that changes in hippocampal microstructure induced by IVMA would be more persistent than those induced by VMA (Sagi et al., 2012). Finally, we explored the interaction between the hippocampus and the motor system in supporting motor learning.

## RESULTS

### VMA and IVMA induce structural changes in the human hippocampus consistent with neuroplasticity

Traditionally, the human hippocampus has been associated primarily with declarative memory. However, recent studies from our group and others have reported its involvement in motor learning, suggesting an expanded role beyond its canonical function to non-declarative, procedural memory (Albouy et al., 2013; Döhring et al., 2017; Long et al., 2018; Schapiro et al., 2019; Jacobacci et al., 2020a; Deleglise et al., 2023). Notably, the bulk of the evidence linking the hippocampus to motor learning, including our own (Jacobacci et al., 2020a), originates from studies using the motor sequence learning task, which depends initially on declarative learning as participants must first memorize the sequence (Robertson et al., 2004; Krakauer et al., 2019). Thus, a fundamental question in neuroscience is to discern whether the participation of the hippocampus in motor learning relates to its canonical declarative role or to processing a general aspect of learning that transcends the declarative/non-declarative distinction.

To address this question, we turned to the classic VMA paradigm, which, unlike MSL, can be easily modified to eliminate the explicit component. This paradigm (Figure 1A, top) consists in making straight pointing movements to visual targets under the influence of an optical rotation applied to the cursor representing the hand (here, 40-degree clockwise -CW), which distorts visuomotor coordination (e.g., Villalta et al., 2015; Solano et al., 2022). Two primary learning processes contribute to VMA, a fast explicit learning process that involves compensating for the optical rotation using a conscious aiming strategy, and a slow implicit learning process involving automatic recalibration (Taylor et al., 2014; Hadjiosif and Krakauer 2021; Tsay et al., 2022; Albert et al., 2022). The former is thought to be driven by errors in achieving the movement goal (task error), whereas, the latter, by errors in predicting the sensory outcome of a movement (sensory-prediction error). Previous studies have demonstrated that explicit learning can be eliminated by clamping the cursor to a fixed movement direction, thereby making task errors constant (Taylor et al., 2014; Morehead et al., 2017). This decoupling forces learning to rely solely on sensory prediction errors, resulting in a fully implicit version of the paradigm (IVMA) (Figure 1A, bottom).

**Figure 1.**
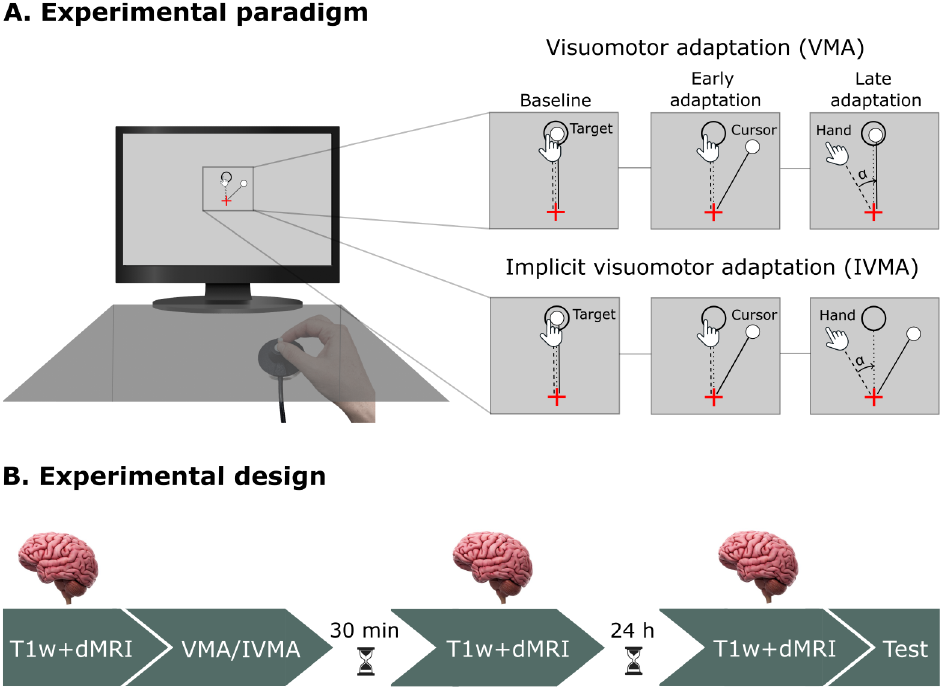
Experimental paradigms and Experimental design. **A)** Experimental paradigms. Subjects trained on one of the two visuomotor adaptation paradigms, consisting of making center-out movements to one of eight visual targets using a joystick while the vision of their hand was occluded. The inset illustrates the visual display of the computer screen across different stages of learning: baseline, early, and late adaptation to an imposed visual perturbation. In the classic paradigm known as visuomotor adaptation (VMA), the cursor represents the movement of the hand in direction and speed. Subjects first performed a set of null trials for familiarization with no perturbation (baseline). Then, an abrupt visual perturbation in the form of an optical rotation (40-degree clockwise) was applied to the cursor representing the hand (early adaptation). With time, subjects learned to compensate for the imposed visual rotation (late adaptation) as indicated by the pointing angle (**α**; i.e., the angle delimited by the straight line connecting the start point with the target position, and the movement direction of the hand). In the implicit visuomotor adaptation (IVMA) task, after the baseline period, the cursor’s trajectory was clamped to 40 degrees clockwise relative to the target’s location throughout the experiment (early adaptation). In this manipulation the cursor matched the speed but not the movement direction of the hand. Decoupling the cursor from the hand’s movement direction avoids improvements in performance based on explicit strategy, forcing subjects to rely on the implicit system. Despite not seeing their trajectories, subjects learned to compensate for the optical rotation (late adaptation). **B)** Experimental Design. Two groups of subjects participated in this study. One group (n = 18) trained on the VMA task, while the other group (n = 22) trained on the IVMA paradigm. Diffusion MRI (dMRI) - and T1 weighted (T1w) images-were acquired before (baseline), 30 min, and 24 h after learning to track the dynamics of changes in microstructure induced by VMA and IVMA in the hippocampus and key motor regions. Overnight memory retention (Test) was assessed using two error-clamp trials. The test session took place after the MRI acquisition obtained 24 h post-training.

Two groups of participants trained on either VMA (n = 18) or IVMA (n = 22), and we examined the impact of motor learning on the temporal dynamics of mean diffusivity (MD) in both the short and long term using diffusion tensor imaging (Figure 1B). Diffusion MRI images were acquired at baseline, 30 min, and 24 h post-training. Evidence from our group and others suggests that sensorimotor memories consolidate within a 4-to 6-hour window (Walker et al., 2003; Lerner et al., 2020; Solano et al., 2024). Thus, our longitudinal design allowed us to sample brain activity early during consolidation and after memory stabilization (Lerner et al., 2020; Solano et al., 2024).

Both groups achieved asymptotic performance after practice on the VMA and IVMA tasks, as reflected by the pointing angle (Figures 2A and 2B, top), and showed similar levels of overnight memory retention assessed using error-clamp trials (VMA, mean ± standard error [SE]: 52.18 ± 4.33%; IVMA: 49.43 ± 4.79% relative to the asymptote).

**Figure 2.**
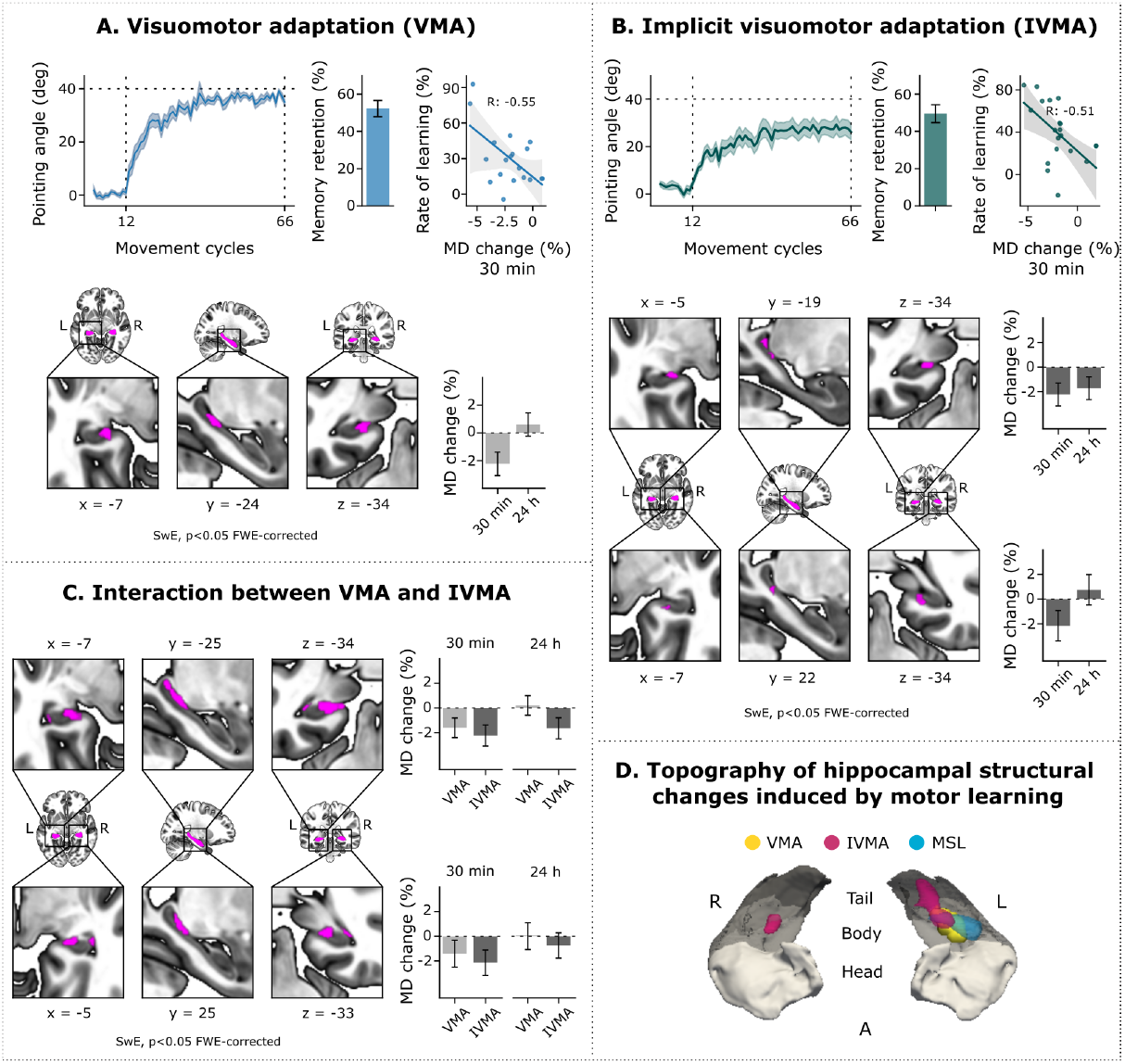
VMA and IVMA induce structural changes in the human hippocampus consistent with neuroplasticity. **A)** Shown are the VMA learning curve (leftmost plot), depicting the pointing angle (mean ± SE) as a function of movement cycles (1 cycle = 8 trials), and memory retention (middle plot) assessed 24 h post-training using error-clamp trials (barplots, mean ± SE relative to the asymptote). The bottom figure depicts the results of applying the SwE statistical model on MD within a bilateral hippocampal mask (depicted in pink in the brain sections) across the three dMRI sessions; the corresponding barplots quantify the % change of MD relative to the baseline ± 95% confidence intervals (CI) for the identified cluster at 30 min and 24 h post-training. VMA induced a significant decrease in MD over the LH 30 min post-learning that reverted to baseline levels by 24 h (p<0.05 FWE-corrected). The rightmost plot illustrates the Pearson correlation between the rate of learning and the percent decrease in MD observed 30 min post-training (r=-0.55, p=0.018). **B)** Shown are the learning curve for IVMA (leftmost plot), and memory retention (middle plot) assessed 24 h post-training using clamp-to-0-degree trials. IVMA induced a significant decrease in MD (bottom figure) over the left (top inset) and right (bottom inset) hippocampi 30 min post-learning; however, only structural changes in the LH persisted 24 h post-training (SwE, p<0.05 FWE-corrected). The rightmost plot shows the Pearson correlation between the rate of learning and the percent decrease in MD observed over the LH 30 min post-training (r=-0.51, p=0.025). **C)** A task x session interaction confirmed the different temporal dynamics of MD for VMA and IVMA (SwE, p<0.05 FWE-corrected; LH: top inset; RH: bottom inset). **D)** Hippocampal clusters identified for VMA and IVMA are rendered in a cartoon of a bilateral 3D hippocampal mask, segmented along its longitudinal axis into head, body, and tail (Iglesias et al., 2015). Structural changes induced by VMA are illustrated in yellow while those induced by IVMA, are in pink. The results from our previous study showing structural changes induced by MSL are overlaid in blue. RH: right hippocampus; LH: left hippocampus. SwE: Sandwich Estimator.

Building on our MSL findings (Jacobacci et al., 2020a), we initially conducted a voxelwise region-of-interest analysis of MD within a bilateral hippocampal mask to identify clusters modulated by motor learning. In line with our previous study, we found that VMA induced a reduction in MD over the left hippocampus 30 min post-learning, which reverted to baseline levels by 24 h (SwE, F(2,17): 15.84, p = 0.001) (Figure 2A, bottom). In contrast, training on IVMA resulted in a reduction in MD over the same area that persisted overnight, i.e., 24 h post-learning (SwE, F(2,21): 20.61, p = 0.001) (Figure 2B, bottom). Notably, IVMA also induced a transient but more subtle reduction in MD over the right hippocampus, which reverted to baseline by 24 h (SwE, F(2,21): 12.29, p = 0.017) (Figure 2B, bottom). The differing temporal dynamics of MD observed for VMA and IVMA are confirmed by the group analysis results shown in Figure 2C (Task x session interaction, SwE, left hippocampus: F(1,34): 7.74, p = 0.001; right hippocampus: F(1,34): 8.91, p = 0.001).

Interestingly, the magnitude of the reduction in MD observed 30 min post-training correlated with the rate of adaptation similarly in both groups (VMA: Pearson correlation, R:-0.55, p=0.018, Figure 2A, IVMA: Pearson correlation, R:-0.51, p=0.025, Figure 2B), indicating that changes in hippocampal microstructure were specific to learning.

Finally, to examine the topography of structural changes induced by motor learning we overlaid the results obtained from training on VMA, IVMA, and MSL (from our previous study) on the bilateral hippocampal mask segmented into head, body, and tail (Iglesias et al., 2015). As depicted in Figure 2D, we found that the three motor learning paradigms induced consistent changes over the posterior portion of the left hippocampus, with a clear anatomical overlap over the hippocampal body. Notably, structural changes induced by IVMA were more extensive, spanning both the body and tail of the left and right hippocampi. Interestingly, neither of the motor tasks recruited the anterior subfield, suggesting a consistent involvement of the posterior hippocampus in procedural motor learning.

Collectively, our results provide strong evidence supporting the role of the hippocampus in implicit motor learning. The overlapping spatial topography of MD changes observed across motor paradigms points to a fundamental involvement of the left posterior hippocampus in procedural motor learning. Contrary to the prediction based on the hippocampus’s canonical role in declarative learning, the suppression of explicit strategies in IVMA resulted in more enduring structural changes. This finding is consistent with the biological implications of a computational model proposing that implicit and explicit learning compete for the available error (Albert et al., 2022), potentially impacting on the neural substrates of memory consolidation.

### During adaptation motor and limbic regions operate together as part of the same network

Abundant experimental evidence underscores the involvement of the cerebellum, the posterior parietal cortex (PPC), and the primary motor cortex as key motor regions involved in visuomotor adaptation. Specifically, it has been shown that the cerebellum and PPC are recruited during early and late phases of acquisition, respectively (Della-Maggiore et al., 2004; Galea et al., 2011; Tseng et al., 2007; Criscimagna-Hemminger et al., 2010; Izawa et al., 2012), whereas the primary motor cortex has been linked to memory consolidation and storage (Richardson et al., 2006; Hadipour-Niktarash et al., 2007; Galea et al., 2011; Orban de Xivry et al., 2011; Landi et al., 2011; Villalta et al., 2015).

Thus, we conducted whole-brain analyses to investigate whether visuomotor adaptation induces structural changes consistent with neuroplasticity in motor regions and whether the dynamics of these changes align with those observed in the hippocampus. We found that VMA induced a transient (Figure 3A) decrease in MD 30 min post-learning over the left lateral cerebellum (lobule VI and Crus I; Grodd et al., 2001; Donchin et al., 2012) and the left PPC, both of which followed the temporal dynamics of MD in the left hippocampus (SwE, p<0.05 FWE-corrected). IVMA also reduced MD in the same network (Figure 3B), but these changes persisted overnight, mirroring the pattern observed in the left hippocampus. In addition, IVMA also induced persistent changes in the contralateral primary motor cortex (Boling et al., 1999) and the right lateral cerebellum, as well as a transient decrease in MD over the right hippocampus (SwE, p<0.05 FWE-corrected). Structural differences across tasks were confirmed by a task-by-session interaction (SwE, p<0.05 FWE-corrected).

**Figure 3.**
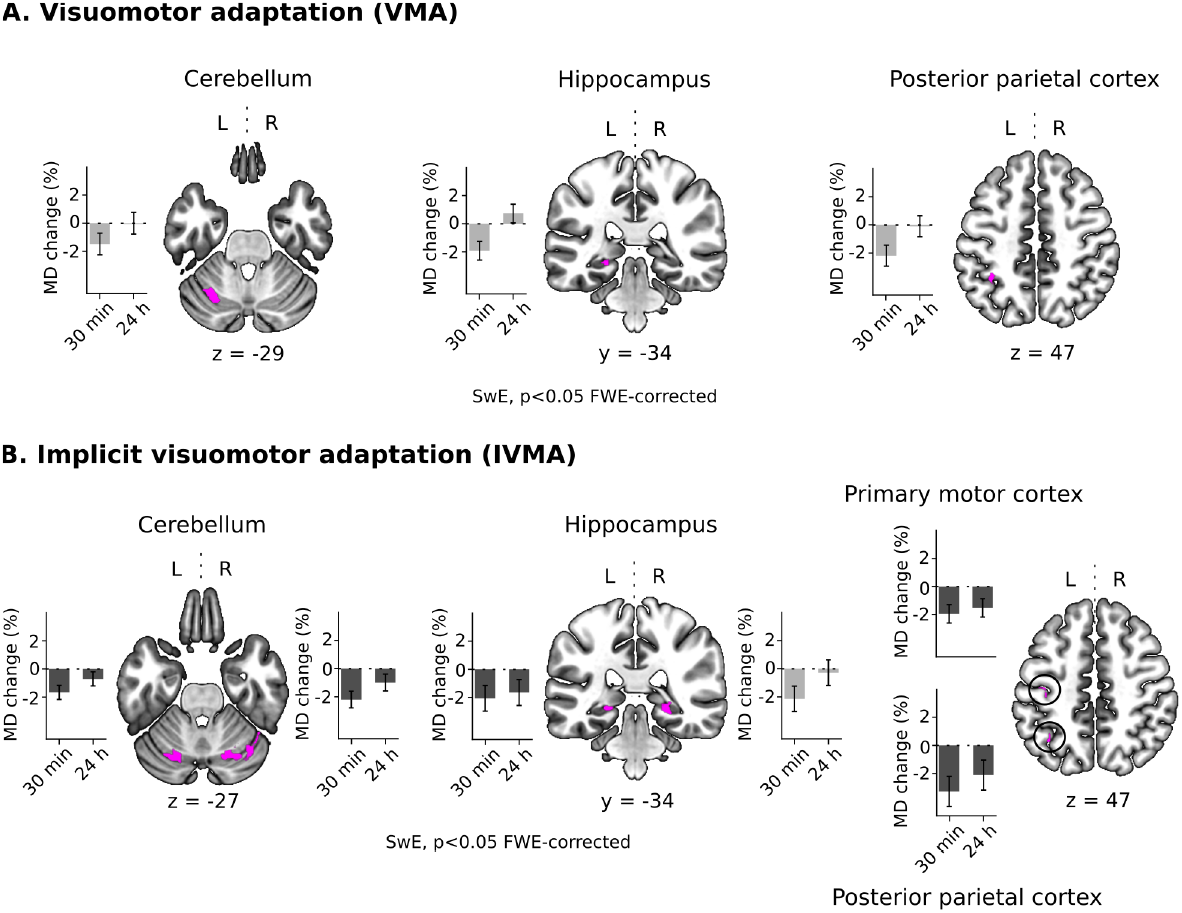
During visuomotor adaptation motor and limbic regions operate together as part of the same network. **A)** SwE conducted across sessions identified structural changes induced by VMA in the left lateral cerebellum, left posterior parietal cortex (PPC), and the same region of the posterior hippocampus detected within the hippocampal mask. In all regions, VMA induced a decrease in MD 30 min post-learning that reverted to baseline values by 24 h. **B)** SwE identified structural changes induced by IVMA over the same brain regions and the primary motor cortex, but in this case, the reduction in MD persisted overnight. IVMA also induced transient structural changes over the right hippocampus. Barplots show the % change ± 95% CI for the time course of MD in these clusters. Transient changes in microstructure are illustrated in light gray while those that persisted overnight are depicted in dark gray.

Collectively, our findings suggest that during visuomotor adaptation, motor and limbic regions function together as part of the same neural network. The distinct temporal dynamics observed across tasks open the possibility that, depending on the reliance on implicit learning, this network may experience either short-or long-lasting microstructural changes. These changes could reflect different biological processes involved in neuroplasticity, ranging from rapid homeostatic adjustments during the initial stages of synaptic potentiation to slower structural remodeling.

Finally, it is important to note that the whole-brain analysis reproduced the anatomical localization and temporal dynamics of hippocampal changes identified with the bilateral mask, further strengthening our findings.

## DISCUSSION

Traditionally, the human hippocampus has been primarily associated with the encoding of declarative memory. In this study, we present compelling evidence for an expanded role in implicit motor learning. Using diffusion MRI to infer structural plasticity, we showed that both the classic visuomotor adaptation paradigm -involving explicit and implicit processing- and an implicit-only version, engaged the left posterior hippocampus in a learning-specific manner. Notably, while the classic paradigm induced transient changes in microstructure, implicit adaptation led to persistent structural alterations. These distinct temporal dynamics of hippocampal plasticity were mirrored by key motor regions involved in adaptation pointing to a close interaction between memory systems. Collectively, our work provides compelling evidence supporting an active role of the hippocampus in procedural motor learning.

In the last decade, functional imaging and neuropsychological studies have reported the participation of the human hippocampus in motor skill learning, particularly within the domain of motor sequence learning (Fletcher et al., 2005; Albouy et al., 2013; Döhring et al., 2017; Long et al., 2018; Schapiro et al., 2019; Jacobacci et al., 2020a; Deleglise et al., 2023). Given that MSL initially involves consciously memorizing the sequence, it is possible that the participation of the hippocampus in motor learning relates to its canonical declarative role. Our results demonstrating the involvement of this structure in implicit visuomotor adaptation are against this possibility, and instead point to a more general function of this structure in memory encoding. Our work aligns with contemporary perspectives from human and non-human studies positing a unifying role of the hippocampus in sequence generation, a fundamental feature inherent in most motor learning paradigms (Buzsáki and Tingley, 2018; Rueckeman et al., 2021; Schwartenbeck et al., 2023).

Notably, changes in microstructure induced by motor sequence learning described in our previous study (Jacobacci et al., 2020a), and visuomotor adaptation reported herein, were consistently localized to the posterior portion of the left hippocampus. The left-lateralization pattern may reflect the asymmetric control of the motor system in planning skilled and reaching movements (De Renzi and Lucchelli, 1988; Schaefer et al., 2007; Yang et al., 2024; Merrick et al., 2022). This hypothesis aligns with the observation that both motor and limbic structures were modulated to a similar extent in both tasks. But how are the motor system and the posterior hippocampus connected? Anatomical evidence indicates that the anterior and posterior hippocampus form part of two distinct networks associated with processing the “what” and “where” aspects of stimuli, respectively (Rolls et al., 2023). Specifically, the posterior portion of the hippocampus is connected with the retrosplenial cortex, and the precuneus, which in turn, connects with the PPC involved in motor planning and sensorimotor adaptation (Della-Maggiore et al., 2004; Kahn et al., 2008; Margulies et al., 2009; Vesia et al., 2010; Adnan et al., 2016, Dalton et al., 2022). Thus, we propose that the consistent topography of hippocampal structural changes observed across motor paradigms may result from the connectivity between limbic and motor systems.

Interestingly, implicit adaptation resulted in more persistent changes in MD than the classic visuomotor adaptation task. This finding can be interpreted in light of the competition model, which proposes that implicit and explicit learning processes compete for error correction during visuomotor adaptation (Albert et al., 2022; Tsay et al., 2022). According to this model, an increase in the explicit system’s response would reduce the extent of implicit adaptation. While we did not assess this directly in VMA, the experimental manipulation implemented here likely led to a greater extent of implicit adaptation compared to the classic paradigm (Morehead et al., 2017). Therefore, we can speculate that the stronger reliance of IVMA on the implicit system may have resulted in a more persistent reduction in MD within the network involving the left hippocampus and key motor regions. Overall, our results suggest that, in the context of visuomotor adaptation, lasting changes in MD are contingent on the relative dominance of the implicit learning component.

What might the differing temporal dynamics in MD reflect at the biological level? Similar to MSL (Jacobacci et al., 2020a), VMA resulted in a transient decrease in this metric, while IVMA led to structural changes that persisted overnight. Rodent studies suggest that the main source of the decreases in MD at the macroscopic MRI resolution is the expansion and/or remodeling of astrocytes (Blumenfeld-Katzir et al., 2011; Sagi et al., 2012; Hofstetter and Assaf, 2017). This may be triggered by various biological phenomena, including acute changes in neuronal activity (Darquié et al., 2001), synaptic potentiation (Sagi et al., 2012), or synapse regulation and stabilization (Kleim et al., 2007; Stevenson et al., 2021). Given that dMRIs were acquired offline the rapid MD alteration observed 30 min post-training is unlikely to reflect increased neuronal activity during learning. Instead, it may result from homeostatic processes such as astroglial swelling, crucial for maintaining the ionic balance during LTP-like plasticity (Sagi et al., 2012). In contrast, the enduring changes induced by IVMA may indicate qualitatively different processes involving fiber organization, such as increased astroglial complexity associated with the stabilization of potentiated and/or new synapses (Kleim et al., 2007; Stevenson et al., 2021).

Finally, our study presents both strengths and limitations. While we would have preferred to combine dMRI with fMRI as in our previous work, this approach is less feasible for visuomotor adaptation. Unlike motor sequence learning, where performance gains occur during the rest periods interspersed with practice, VMA improvements take place during active practice. Moreover, during initial practice in VMA, increases in activity may result from a mix of factors including changes in movement kinematic, error-driven processes, and force (Diedrichsen et al., 2005) that may mask and/or confound learning-related changes in hippocampal activity (Jacobacci et al., 2020a). While our findings rely mostly on microstructure, several facts confirm the specificity and reliability of our work. First, rapid MD changes induced by MSL and visuomotor adaptation tasks related to the rate of learning, pointing to the hippocampal involvement in the encoding/early consolidation of sensorimotor memories. Second, all three tasks induced structural changes that converged to the posterior portion of the left hippocampus supporting the generalizability of our results to procedural motor learning. Third, the whole-brain analysis identified regions of the motor system that are critically involved in visuomotor adaptation such as the motor cortex, the PPC and the cerebellum (Della-Maggiore et al., 2004; Tseng et al., 2007; Chrishimanga-Hemminger et al., 2010).

In conclusion, our study underscores the specific engagement of the hippocampus in implicit motor learning. The convergence of structural changes across both visuomotor and motor sequence learning paradigms points to a broader role of the hippocampus in procedural motor learning. Moreover, all motor paradigms recruited the left posterior hippocampus in temporal conjunction with key motor regions, a finding in line with the left-lateralized control of movement planning. Finally, our results suggest that in visuomotor adaptation, persistent changes in microstructure depend on the reliance on the implicit learning component. Our work points to the existence of common mechanisms in memory encoding across different domains regardless of the explicit or implicit nature of the task.

## ACKNOWLEDGMENTS

We thank the Argentinian Agency for the Promotion of Science and Technology (FONCyT: PICT2018-1150 & PICT2019-2156) for their financial support.

## METHODS

### Participants

A total of forty participants aged between 18 and 34 years old (mean ± SD = 24.8 ± 3.9 years old; 26 females) were enrolled in the study. All participants were healthy volunteers with no self-reported history of psychiatric, neurological, or cognitive disorders, nor any history of sleep disturbances. Participants were instructed to abstain from alcohol on the day before and during the experiment, as well as to maintain their regular sleep habits. All subjects were right-handed as assessed by the Edinburgh handedness inventory (Oldfield 1971). Written consent was obtained from all participants, and they were remunerated for their time and inconvenience. The experimental procedure was approved by the local Ethics Committee (University of Buenos Aires) and was conducted following the Declaration of Helsinki.

### Experimental paradigms

#### Visuomotor adaptation (VMA)

The VMA paradigm used here has been described in previous studies (e.g., Villalta et al. 2015; Lerner et al. 2020; Solano et al. 2022) and is summarized here. This center-out task requires participants to hit one of eight visual targets displayed concentrically around the start position and equidistantly, on a computer screen using a cursor controlled with the thumb and index finger of their right-dominant hand (Figure 1A). In this paradigm, the vision of the hand is occluded. On each trial, subjects executed a shooting movement to one of the eight targets presented individually on the screen in a pseudo-randomized order. One cycle consisted of eight trials directed to each target; 11 cycles composed a block. Continuous visual feedback on the hand’s position was provided through the cursor from the onset of each trial until the movement was completed. To avoid online corrections that would lead to submovements, the joystick’s gain was set to 1.4, so that a displacement of 1 cm of the tip of the joystick moved the cursor on the screen by 1.4 cm. According to previous pilot data from our lab, this gain yields straight paths with little or no online corrections (Villalta et al., 2015).

The VMA paradigm involved three different types of trials. During null trials, the cursor movement directly mapped the joystick’s movement in native coordinates. During perturbed trials, a CW 40-degree optical rotation was imposed on the cursor, deviating its trajectory. During error-clamp (EC) trials, the cursor trajectory was artificially manipulated to simulate “straight” paths to the target, resembling that of correct trials, by projecting the actual cursor’s movement onto a straight line with additional variability (0 ± 10 degrees, mean ± standard deviation). EC trials allowed estimating memory retention by eliminating learning from error (Criscimagna-Hemminger and Shadmehr, 2008).

#### Implicit visuomotor adaptation (IVMA)

Two main learning processes contribute to visuomotor adaptation, a fast explicit learning process that involves compensating the optical rotation using a conscious aiming strategy, and a slow implicit learning process involving automatic recalibration (Taylor et al., 2014; Tsay et al., 2022; Albert et al., 2022). The former is thought to be driven by errors in achieving the movement goal (task error), whereas, the latter, by errors in predicting the sensory outcome of a movement (sensory prediction error). To study implicit adaptation in isolation of the explicit component, we fixed task error by clamping the cursor direction to 40 degrees CW relative to the target’s position throughout training (Figure 1A, bottom). In this manipulation the cursor matches the speed of the hand while maintaining no contingency with the hand angle, ensuring that learning is driven by the implicit component (sensory prediction error) (Morehead et al. 2017). Despite their awareness of the manipulation, participants adapt to the perturbation implicitly (Kim et al. 2018). We refer to this task as implicit visuomotor adaptation (IVMA).

As in the VMA paradigm, each cycle consisted of 8 trials and each block consisted of 11 cycles. The IVMA paradigm involved three different types of trials. During null trials, the cursor movement directly mapped the joystick’s movement. During clamped-to-40-degree trials, a fixed CW 40-degree visual rotation was imposed on the cursor relative to the visual target’s position. Finally, during clamped-to-0-degree trials, the cursor trajectory was fixed to the visual target position. The latter was used to quantify memory retention in this paradigm because it avoids further learning while providing no veridical feedback on hand movement.

Both VMA and IVMA tasks were implemented in MATLAB (The MathWorks, Inc.) using the Psychophysics Toolbox v3 (Brainard 1997).

### Experimental design and procedure

This study was designed to examine the involvement of the hippocampus in implicit motor learning. To this aim, we used MD, an indirect marker of structural plasticity, to quantify changes in microstructure induced by the classic VMA paradigm –relying both on explicit and implicit learning–, and an implicit-only version (IVMA), in the short and the long term. To be able to contrast the visuomotor adaptation tasks with the motor sequence learning task used in our previous study (Jacobacci et al., 2020a), T1w and dMRI images were acquired following the same longitudinal design: before, 30 min, and 24 h after learning (Figure 1B). Overnight memory retention was assessed 24 h after training.

Two groups of subjects participated in this study. One group (n = 18; mean ± SD = 25.3 ± 4.2 years old; 12 females) trained on the VMA task, while the other group (n = 22; mean ± SD = 24.7 ± 3.8 years old; 14 females) trained on the IVMA paradigm. Both groups were instructed to perform a shooting movement to one of the eight targets with the cursor, as soon as it appeared on the screen. Before practice, subjects became familiar with the setup by performing a few center-out movements with veridical feedback during null trials. Additionally, participants in the IVMA group were told of the clamped-cursor manipulation and were instructed to ignore the cursor direction and aim directly at the target throughout the experiment. Participants in the VMA group performed 1 block of null trials followed by 6 blocks of perturbed trials, while participants in the IVMA group performed 1 block of clamped-to-0-degree trials followed by 6 blocks of clamped-to-40-degree trials. Training for both tasks lasted approximately 25 minutes.

### Data acquisition and preprocessing

#### Magnetic resonance (MR) images

MR images were acquired using a 3T Siemens Tim TRIO MRI scanner located at the Instituto Angel Roffo, University of Buenos Aires (Argentina), equipped with a 12-channel head coil. MRI images were acquired using a longitudinal design before (baseline), 30 min, and 24 h post-training on VMA or IVMA, as indicated in Figure 1B. On each session, an anatomical T1-weighted MPRAGE image was collected with the following parameters: repetition time (TR) = 2530 ms; echo time (TE) = 2.17 ms; inversion time (TI) = 1100 ms; flip angle (FA) = 7°; bandwidth (BW) = 199 Hz/Px; field of view (FOV)= 256 × 256 mm^2^; acquisition matrix = 256 × 256; 1-mm isotropic voxels; slices = 176; parallel acquisition = GRAPPA mode, acceleration factor= 2. The acquisition was performed in the sagittal plane.

Three dMRI acquisitions were collected on each session using the SE-EPI sequence with the following parameters: two dMRIs were acquired with an anterior-posterior (A-P) phase encoding direction and one with the opposite direction (P-A); multiband acceleration factor (Xu et al., 2013; Uğurbil et al., 2013) = 2; TR = 5660 ms; TE = 89 ms; FA = 90°; BW = 1488 Hz/Px; slices = 76; 2-mm isotropic voxels; FOV = 240×240 mm^2^; EPI factor = 120; acquisition: interleaved; 30 monopolar gradient directions with b-value = 1000 s/mm^2^. Furthermore, seven b0s were acquired using the same phase encoding direction: two at the beginning of the acquisition, one at the end, and the rest interleaved every five b-1000 volumes.

Preprocessing and normalization of the dMRIs were performed using the FMRIB Software Library (FSL; University of Oxford) (version 6.0.3) and the Advanced Normalization Tools (ANTs; Wellcome Department, UCL) (version 2.4.2). The preprocessing steps were conducted separately for each scanning session and included the correction of susceptibility-induced distortions using FSL’s Topup (Andersson et al., 2003), correction of eddy currents induced distortions, head motion correction, and b-vector rotation using FSL’s eddy tool (Andersson et al., 2016). Subsequently, FSL’s DTIfit was used to fit a diffusion tensor model and produce the scalar maps for FA and MD metrics. Following preprocessing, DTI scalar maps were normalized to MNI152 stereotaxic space using FA as an intermediate template. We used ANTs for normalization in a pipeline created by our group to minimize across-session test-retest reproducibility error (Jacobacci et al., 2020b). Finally, normalized DTI measures were smoothed with FSL’s smoothing tool with a 4-mm full-width at half maximum (FWHM) Gaussian kernel.

#### Hippocampal subfields

Human hippocampal subfields were obtained using Freesurfer (version 7.0). The FSL’s MNI T1 brain image of 1-mm resolution was processed through Freesurfer’s *recon-all* function. Subsequently, we used the *segmentHA_T1* optimized pipeline to extract the hippocampal subfields. This specific pipeline allowed segmenting the hippocampus along its longitudinal axis, into the head, body, and tail (Iglesias et al., 2015).

### Data analysis and statistics

#### Behavior

Motor performance for both VMA and IVMA tasks was quantified based on the pointing angle, that is the angle determined by the movement direction of the joystick and the line segment connecting the start point with the target position (e.g., Lerner et al., 2020; Solano et al., 2022). Trials in which the pointing angle exceeded 120° were excluded from subsequent analysis. Given that trials were organized into cycles, trial-by-trial data were transformed into cycle-by-cycle time series by computing the median pointing angle for each cycle and subject. The asymptotic performance for each participant was determined by calculating the mean of the last five cycles during the final adaptation block.

To assess memory retention for each task and each subject we used two EC cycles for VMA and two clamp-to-0-degree cycles for IVMA, relative to each subject’s asymptotic performance according to this equation: (PA_A_-*abs*(PA_A_-PA_EC/clamp-to-0_))*100/PA_A_, where PA_A_ is the pointing angle of the asymptote and PA_EC/clamp-to-0_is the pointing angle of the two retention cycles. Finally, this percentage measure was averaged across the two cycles. The rate of learning was assessed by calculating the median of the pointing angle across the first four cycles -excluding the first one-during the adaptation phase (Huberdeau et al., 2015; Kim et al., 2018), and expressed as a percentage of the pointing angle asymptote. Three subjects from the IVMA group were excluded from the behavioral analysis because their pointing angle of the asymptote was less than 0 degrees.

#### dMRIs

Longitudinal changes in MD induced by VMA and IVMA across sessions were computed using a non-parametric Sandwich Estimator (SwE, Guillaume et al., 2014) model with a permutation-based threshold-free cluster enhancement (TFCE) approach (Smith and Nichols, 2009). SwE has been specifically designed for accurate modeling of longitudinal and repeated-measures neuroimaging data (version 2.0.0). To detect hippocampal changes in microstructure across sessions (baseline, 30 min, 24 h), statistical analyses were conducted separately for each task within a bilateral hippocampal mask derived from the Harvard-Oxford subcortical structural probability atlas (>30% probability, FSL). To compare changes induced by VMA vs IVMA we conducted a task (VMA, IVMA) by session (baseline, 30 min, 24 h) interaction analysis within the same hippocampal mask. Furthermore, to identify changes in microstructure at the level of the motor system we conducted a separate whole-brain SwE analysis across sessions as implemented in our previous study (Jacobacci et al., 2020a) for VMA and IVMA, and their interaction. All non-parametric SwE analyses were conducted based on 1,000 permutations, and significant clusters were identified using a family-wise error (FWE) corrected p-value < 0.05.

Following cluster identification, MD values were extracted from all voxels within each cluster for each subject and session, and the median MD was computed. MD changes for each cluster at 30 min and 24 h were expressed relative to the baseline (e.g., (30 min MD - baseline MD)/baseline MD). For each cluster and each session, we computed the mean percent change in MD across subjects and determined the 95% confidence intervals. This was achieved using the *summarySEwithin* function from the *Rmisc* package in R (Team, R. C, 2020). This function is designed to normalize within-subject data, effectively by removing between-subject variability using the method proposed by Morey and colleagues (2008).

To anatomically compare structural changes induced by different motor learning tasks, we conducted the interaction between the MSL and the active control condition (CTL task) from our previous study (Jacobacci et al., 2020a), within the bilateral hippocampal mask using the same SwE approach, and overlaid these results from the ones obtained in the current study.

In our prior MSL study, we found that rapid changes in MD related to gains in performance during the early stages of learning. To evaluate if changes in MD induced by VMA and IVMA were also specific to memory encoding we computed a Pearson correlation (R Core Team, 2013) between the percent change in MD observed 30 min post-learning and the rate of learning.

Of note, TFCE parameters for all SwE analyses within the hippocampal mask were set to H=4.5 and E=0.5, to conform to a more conservative approach, whereas, those for the whole-brain analysis, were set to relatively less conservative parameters to ensure good sensitivity (H=4 and E=0.5). Note, however, that the chosen TFCE parameters were always more conservative than the parameter settings recommended by Smith and Nichols (H=2 and E = 0.5).

